# Temporal control of late replication and coordination of origin firing by self-stabilizing Rif1-PP1 hubs in *Drosophila*

**DOI:** 10.1101/2022.01.14.476400

**Authors:** Chun-Yi Cho, Charles A. Seller, Patrick H. O’Farrell

**Affiliations:** Department of Biochemistry and Biophysics, University of California, San Francisco, San Francisco, California 94158, USA

**Keywords:** Rif1, protein phosphatase 1, DDK, DNA replication timing, *Drosophila*

## Abstract

In the metazoan S phase, coordinated firing of clusters of origins replicates different parts of the genome in a temporal program. Despite advances, neither the mechanism controlling timing nor that coordinating firing of multiple origins is fully understood. Rif1, an evolutionarily conserved inhibitor of DNA replication, recruits protein phosphatase 1 (PP1) and counteracts firing of origins by S-phase kinases. During the mid-blastula transition (MBT) in *Drosophila* embryos, Rif1 forms subnuclear hubs at each of the large blocks of satellite sequences and delays their replication. Each Rif1 hub disperses abruptly just prior to the replication of the associated satellite sequences. Here, we show that the level of activity of the S-phase kinase, DDK, accelerated this dispersal program, and that the level of Rif1-recruited PP1 retarded it. Further, Rif1-recruited PP1 supported chromatin association of nearby Rif1. This influence of nearby Rif1 can create a “community effect” counteracting kinase-induced dissociation such that an entire hub of Rif1 undergoes switch-like dispersal at characteristic times that shift in response to the balance of Rif1-PP1 and DDK activities. We propose a model in which the spatiotemporal program of late replication in the MBT embryo is controlled by self-stabilizing Rif1-PP1 hubs, whose abrupt dispersal synchronizes firing of associated late origins.

**SIGNIFICANCE:** Seventy years ago, it was discovered that large domains of the eukaryotic genome replicate at different times. Detailed descriptions left significant questions unresolved. How are the many origins in the large domains coordinated to fire in unison? What distinguishes different domains and gives rise to a temporal program? When *Drosophila* embryos first establish late replication, an inhibitor of DNA replication, Rif1, forms hubs over domains of late replicating DNA. Rif1 recruits protein phosphatase 1 (PP1), which prevents kinases from dispersing Rif1 hubs or activating associated origins. When kinase activity eventually exceeds a hub-specific threshold, the self-stabilization of Rif1-PP1 breaks down, hubs disperse abruptly, and all associated origins are free to initiate.

## INTRODUCTION

During a typical metazoan cell cycle, large genomic domains initiate their replication at distinct times in S phase (1). Cytological studies over 60 years ago revealed that DNA sequences in the compacted heterochromatin replicate later in S phase compared to euchromatin (2, 3). These early studies and recent detailed analyses revealed a complex program among “late” replicating domains, in which different domains initiate replication with a specific delay (4). Execution of this stereotyped schedule occupies much of the S phase and must finish before mitosis. Despite recent advances in genomic methods for profiling global replication timing (5), the basis of the timing control is not yet solved, and we do not know how multiple origins are coordinated to fire together especially within repetitive DNA sequences.

The *Drosophila* embryo offers a unique setting in which to examine the control of temporal programing of replication. In the earliest nuclear division cycles, there is no late replication—closely spaced origins throughout the genome initiate replication rapidly at the beginning of interphase, and their simultaneous action results in an extraordinarily short S phase (3.5 min) (6, 7). Late replication is developmentally introduced during the synchronous blastoderm nuclear division cycles, first influencing pericentric satellite sequences that form a major part of metazoan genomes (over 30% in *Drosophila*) (8). Individual blocks of satellite DNA are typically several megabase pairs in length, each composed of a different simple repetitive sequence. During the 14^th^ cell cycle at the mid-blastula transition (MBT), the ~6000 cells of the entire embryo progress synchronously through a temporal program in which the different satellites are replicated with distinctive delays (4), dramatically extending the duration of S phase.

The initial onset of late replication during development provides a simplified context in which to define its mechanism, because numerous complex features associated with replication timing have not yet been introduced. For example, chromatin states can have major impacts on replication timing. Consistent with this, late-replicating satellite sequences are usually heterochromatic, carrying the canonical molecular marks of constitutive heterochromatin (histone H3 lysine 9 methylation and HP1). During initial *Drosophila* embryogenesis, the satellites lack significant levels of these marks, and they replicate in sync with the rest of the genome (4, 9). Surprisingly, the introduction of the delays in replication to the satellite sequences precedes a major wave of heterochromatin maturation in the blastoderm embryo (4, 9–11). Furthermore, in a *Rif1* null mutant, the S phase of cycle 14 is significantly shorter, and the late replication of satellite sequences is largely absent even though HP1 recruitment appears normal (10). Thus, a Rif1-dependent program bears virtually full responsibility for the S-phase program at the MBT.

Rif1 is a multifunctional protein with an evolutionarily conserved role in regulating global replication timing (17). In species from yeast to mammals, mutation or depletion of Rif1 disrupts genome-wide replication timing (10, 12–16, 18). Studies in a variety of systems revealed several aspects of Rif1 function. Yeast Rif1 associates with late origins (14, 19, 20), while both *Drosophila* and mammalian Rif1 binds broadly within large late-replicating domains (10, 12, 13, 21). Rif1 has a conserved motif for interacting with protein phosphatase 1 (PP1) (22), and the mutations in PP1-interacting motifs lead to hyperphosphorylation of MCM helicase in the pre-replicative complex (pre-RC) and the disruption of global replication timing (16, 23–27). Rif1 itself also harbors many sites recognized by S-phase kinases, including CDK and DDK, near its PP1-interacting motifs. In yeast, both a Rif1 mutant with phosphomimetic changes at these phosphorylation sites and a null mutation of Rif1 partially restore the growth defect of DDK mutants (24, 25). These data suggest an interplay of Rif1 and DDK, wherein DDK acts first upstream of Rif1 phosphorylating it to disrupt its interaction with PP1, thus lowering the threshold of S-phase kinase activities required for origin firing. Second, DDK acts downstream to directly phosphorylate pre-RC and trigger origin firing (28–30). However, how these various features of Rif1 and DDK functions are integrated over large genomic regions to provide a domain-level control of replication timing remains elusive.

Studies in flies indicate that Rif1 has adopted a developmental role in governing the onset of the late replication program described above. During the early embryonic cell cycles, high Cdk1 activity inhibits maternally deposited Rif1 and promotes synchronous firing of origins throughout the whole genome to ensure completion of DNA replication during the short interphases (4, 10, 31). When the embryo enters the MBT in cycle 14, abrupt down-regulation of Cdk1 derepresses Rif1, which then associates with chromatin and accumulates in semi-stable foci (hubs) at satellite DNA loci (4, 10, 31, 32). High-resolution live microscopy reveals that different Rif1 hubs disperse abruptly at distinct times, followed by PCNA recruitment as the underlying sequences replicate (10). Mutated Rif1 that is non-phosphorylatable at a cluster of CDK/DDK sites fails to dissociate from satellite DNA and dominantly blocks the completion of satellite DNA replication before mitosis of cycle 14. Conversely, ectopically increasing CDK activity in cycle 14 shortens the persistence of endogenous Rif1 foci and advances the replication program (10). These findings suggest that each Rif1 hub maintains a local nuclear microenvironment high in Rif1-recruited PP1 that inhibits DNA replication, and that kinase-dependent dispersal of Rif1 hubs is required to initiate the replication of satellite sequences. If we understood what coordinates Rif1 dispersal throughout the large Rif1 hubs, this model could explain how firing of clusters of the underlying origins is coordinated and how replication of different satellites occurs at distinct times. However, the precise mechanisms controlling the dynamics of Rif1 hubs remain unclear.

Since Rif1 can recruit PP1 and form phosphatase-rich domains in the nucleus, we hypothesized that localized PP1 counteracts kinase-induced Rif1 dissociation so that the Rif1 hubs are self-stabilizing. If this self-stabilization is communicated within each hub, a breakdown in self-stabilization would lead to a concerted collapse of the entire hub and allow origin firing throughout the associated satellite sequence. Our findings herein indicate that the opposing actions of phosphatase and kinase combined with communication within the hubs create a switch in which a large phosphatase-rich domain is stable until kinase activity overwhelms the phosphatase. We propose that for large late-replicating regions of the genome, recruitment of Rif1-PP1 creates a new upstream point of DDK-dependent regulation in which DDK triggers the collapse of the phosphatase-rich domain to create a permissive environment for kinase-induced firing of all previously repressed origins.

## RESULTS

### The recruitment of PP1 is required for the formation of stable Rif1 hubs at domains of satellite DNA

To test the hypothesis that Rif1 hubs are stabilized by the action of local PP1, we used CRISPR-Cas9 to generate a Rif1^pp1^-GFP mutant allele by mutating the PP1-docking motif in a GFP-tagged form of the endogenous Rif1 from RVSF to RVSA (Figure 1A), which strongly disrupts the interaction with PP1 (33). Flies homozygous for the Rif1^pp1^-GFP allele are healthy and fertile, as previously reported for an independently generated untagged Rif1^pp1^ allele (34). We then asked how the lack of PP1 recruitment impacts the stability of Rif1 hubs and S-phase progression during nuclear cycle 14 (NC14) by time-lapse confocal microscopy. The wild-type Rif1-GFP forms nuclear foci at satellite DNA upon entering interphase in NC14, and those foci disappear progressively in accordance with the temporal program of late replication (Figure 1B) (10). In contrast, we found that the mutant Rif1^pp1^-GFP only formed a few small foci that rapidly dissociated from chromatin during early NC14 (Figure 1B).

**Figure 1.**
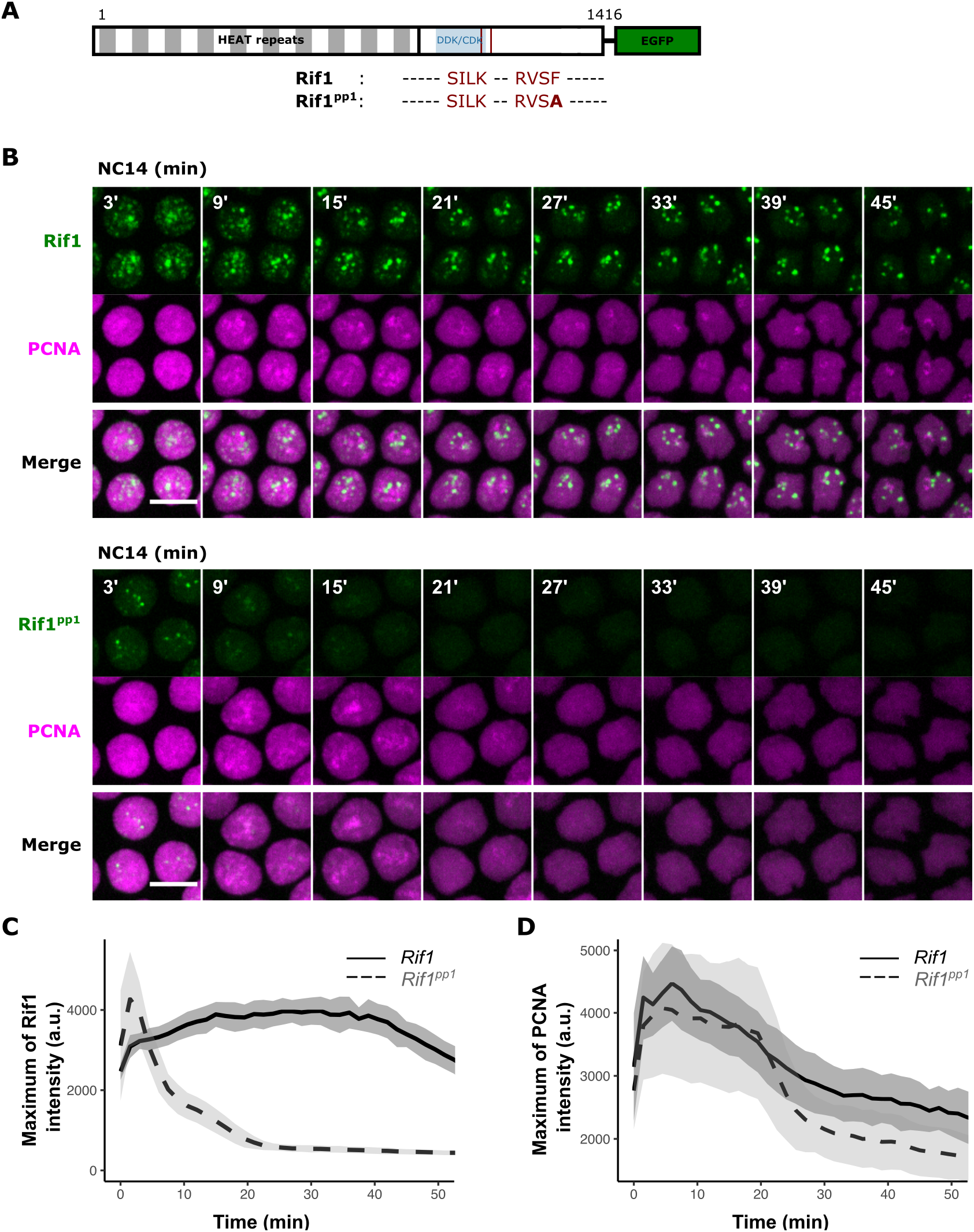
The recruitment of PP1 is required for the formation of stable Rif1 foci at satellite DNA and late replication. (A) Schematic diagram of the *Drosophila* Rif1 protein with wild-type or mutated PP1-interacting motif. Red lines indicate the binding motifs for protein phosphatase 1 (PP1) with amino acid sequences shown below. Striped box indicates the N-terminal region containing 19 HEAT repeats. Blue box indicates region containing 15 putative phosphorylation sites by cyclin-dependent kinase (CDK) and Dbf4-dependent kinase (DDK). (B) Time-lapse imaging of endogenously GFP-tagged Rif1 or Rif1^pp1^ along with mCherry-PCNA in embryos during nuclear cycle 14 (NC14) at the MBT. The start of interphase is set as time point 0’. Scale bars, 6 μm. (C) Maximum of GFP-tagged Rif1 or Rif1^pp1^ fluorescent intensity in the nucleus during NC14. Data are mean ± s.d. (n = 9 embryos). (D) Maximum of mCherry-PCNA fluorescent intensity in the nucleus during NC14. Data are mean ± s.d. (n = 9 embryos). a.u., arbitrary unit.

To compare the dynamics of wild-type and mutant Rif1 foci, we quantified the maximal nuclear intensity of GFP-tagged Rif1 or Rif1^pp1^ as a proxy for their chromatin association over the course of NC14. Indeed, although the initial peak fluorescent intensity at the few Rif1^pp1^ foci was comparable to that of the more abundant and larger Rif1 foci, it quickly declined and diminished to nuclear background around 20 minutes in NC14 (Figure 1C). In contrast, the fluorescent intensity of wild-type Rif1 foci persisted at high level for over 50 minutes, indicating their stable accumulation at satellite DNA. These results support the essential role of PP1 in stabilizing Rif1 foci at late-replicating domains.

To examine the impact of precocious Rif1 foci dissociation on S-phase progression, we performed simultaneous live imaging of mCherry-PCNA, which is recruited to replication forks and marks sites of active DNA replication (35). In the wild-type Rif1-GFP embryo, waves of late-replicating foci marked by bright PCNA signal could be observed throughout the majority of NC14 (Figure 1B). In the mutant Rif1^pp1^ embryo, PCNA foci disappeared in the nucleus precociously around 30 minutes in NC14 and never reappeared, indicating the corresponding shortening of S phase due to the destabilization of Rif1^pp1^ foci (Figures 1B and 1D). We conclude that the ability of Rif1 to recruit PP1 contributes to the recruitment of Rif1 to satellites and to their stable association, as well as contributing to the delayed replication of satellite sequences and the overall duration of S phase.

By the end of NC14 and entry into cycle 15, most satellite sequences have formed more mature heterochromatin and have clustered together as a coherent chromocenter (4, 9). We asked whether stable Rif1 recruitment to this more mature heterochromatin also depends on its PP1-interacting motif. The wild-type Rif1-GFP embryos entered cycle 15 with a compacted chromocenter that rapidly recruited Rif1, and the replication of the chromocenter was delayed and coupled with the progressive loss of Rif1 in mid-to-late S phase 15 (Figure S1). In contrast, the recruitment of Rif1^pp1^-GFP was reduced, residual foci dispersed precociously, and early emergence of foci of PCNA indicated precocious heterochromatin replication (Figure S1). Therefore, Rif1-recruited PP1 contributes to the formation of stable Rif1 hubs at satellite DNA packaged into constitutive heterochromatin and to the timing of its eventual replication.

### Rif1-recruited PP1 acts in *trans* to stabilize Rif1 hubs

PP1 might only promote the chromatin association of Rif1 to which it is recruited, or Rif1-recruited PP1 might act in *trans* on neighboring Rif1 within the hub. We thus wanted to determine whether the defects in Rif1^pp1^ recruitment and stability (Figure 2A, upper panel) could be rescued by the presence of wild-type Rif1. In the heterozygous Rif1^pp1^-GFP/+ embryos, both the initial recruitment and stability of Rif1^pp1^-GFP foci were significantly rescued (Figure 2A, lower panel). Furthermore, the eventual disappearance of GFP foci was still followed shortly by the recruitment of PCNA in the heterozygous embryos (Figure S2A), indicating coordinated dissociation of wild-type Rif1 with the tagged Rif1^pp1^. We conclude that Rif1-recruited PP1 can act in *trans* between different Rif1 proteins. This behavior could result from the formation of mixed oligomers of Rif1 and Rif1^pp1^ (36), and/or from an ability of the recruited PP1 to stabilize neighboring Rif1 (action in *trans*) and so produce a “community effect” stabilizing Rif1 binding throughout an individual Rif1 hub (see Discussion).

**Figure 2.**
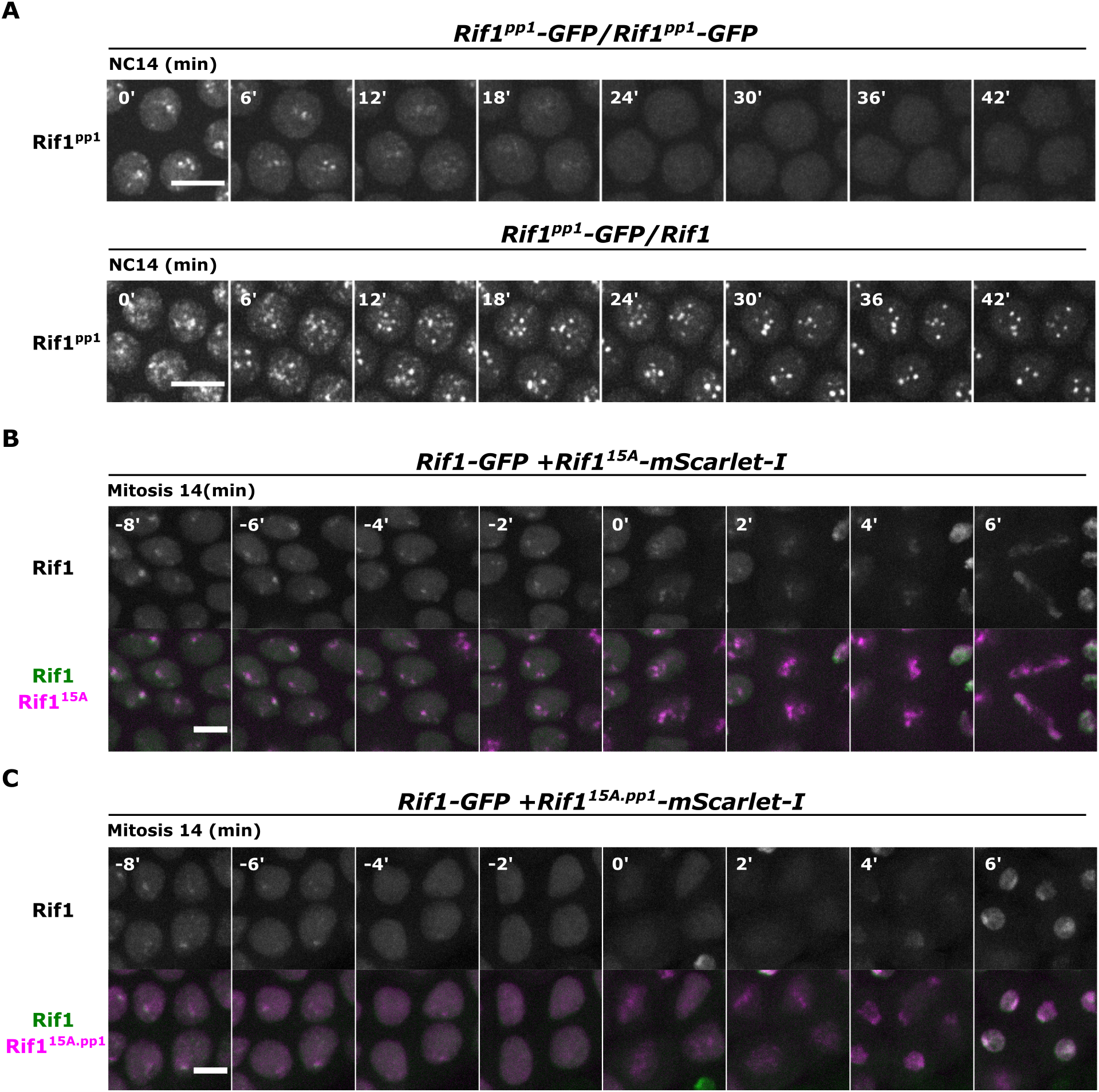
Rif1-associated PP1 can act in *trans* to stabilize Rif1 foci. (A) Time-lapse imaging of Rif1^pp1^-GFP in embryos from homozygous female (top) or from female heterozygous with the wild-type Rif1 allele (bottom) during NC14. The start of interphase is set as time point 0’. Brightness and contrast were independently adjusted for comparison. (B) Time-lapse imaging of endogenous Rif1-GFP and ectopically expressed Rif1^15A^-mScarlet-I lacking 15 CDK/DDK phosphorylation sites (see Figure 1A). Snapshots from late cycle 14 as nuclei went through mitosis are shown. (C) Time-lapse imaging of endogenous Rif1-GFP and ectopically expressed Rif1^15A.pp1^-mScarlet-I lacking 15 CDK/DDK phosphorylation sites and PP1-interacting motif. All scale bars, 6 μm.

To further test the ability of Rif1-associated PP1 to act in *trans* to stabilize Rif1 foci, we asked how ectopic Rif1^15A^, which lacks a cluster of CDK/DDK phosphorylation sites and forms hyperstable foci (10), affects the localization of endogenous Rif1. We injected mRNAs encoding Rif1^15A^ tagged with mScarlet-I into Rif1-GFP embryos before NC13 and performed live imaging through NC14. In support of the hypothesis, we observed that Rif1-GFP colocalized with Rif1^15A^-mScarlet-I foci and that both signals persisted into mitosis, beyond the time of normal dispersal of Rif1-GFP prior to the end of S phase (Figures 2B and S2B). After a prolonged metaphase, cells eventually entered anaphase and produced extensive chromosomal bridges (Figure 2B), as expected from the previously reported ability of Rif1^15A^ to persistently inhibit replication. Thus, stably bound Rif1^15A^ can promote the stable binding of wild-type Rif1.

To further assess whether the recruitment of PP1 contributes to the above outcomes, we repeated the experiment with a Rif1^15A.pp1^ construct, whose PP1-interacting motifs were further mutated to SAAA/RVSA. Interestingly, Rif1^15A.pp1^ was still capable of reinforcing endogenous Rif1-GFP foci in early NC14 (Figure S2B). However, unlike Rif1^15A^, Rif1^15A.pp1^ did not prolong Rif1-GFP association during late interphase or mitosis, and chromosomes segregated normally upon anaphase entry (Figure 2C). Therefore, both the inhibition of replication and the protection of Rif1 foci from kinase-induced dissociation by Rif1^15A^ are dependent on the ability of Rfi1^15A^ to recruit PP1. We conclude that the behavior of Rif1-GFP was non-autonomously influenced by the PP1 recruited by stably bound Rif1^15A^, again supporting the conclusion that Rif1 binding is influenced by its neighbors.

### The concentration of Rif1 determines the timing of its own dissociation from satellite DNA

The self-stabilizing regulation of Rif1 hubs suggests that the dosage of Rif1, and thus recruited PP1, might play an integral role in timing their dissociation from satellite DNA. To test the dosage-dependent stability of Rif1 foci, we asked how reducing the total abundance of Rif1 impacts S-phase progression at the MBT. As most of the Rif1 protein at this stage is maternally provided, we examined embryos from females that were transheterozygous for Rif1-GFP tagged at the endogenous locus paired with either a *rif1* null allele (deletion allele) or non-tagged Rif1 as control. In the control Rif1-GFP/+ embryos, the temporal program of Rif1 foci dissociation played out over 80 min as it did in homozygous Rif1-GFP embryos (Figure S3A). In contrast, in the Rif1-GFP/*rif1* embryos, both the persistence of Rif1 foci and S-phase duration were significantly shortened (Figures 3A and S3B). Thus, reducing the dose of Rif1 destabilized the Rif1 foci at satellite DNA loci and advanced the replication timing of the associated DNA.

**Figure 3.**
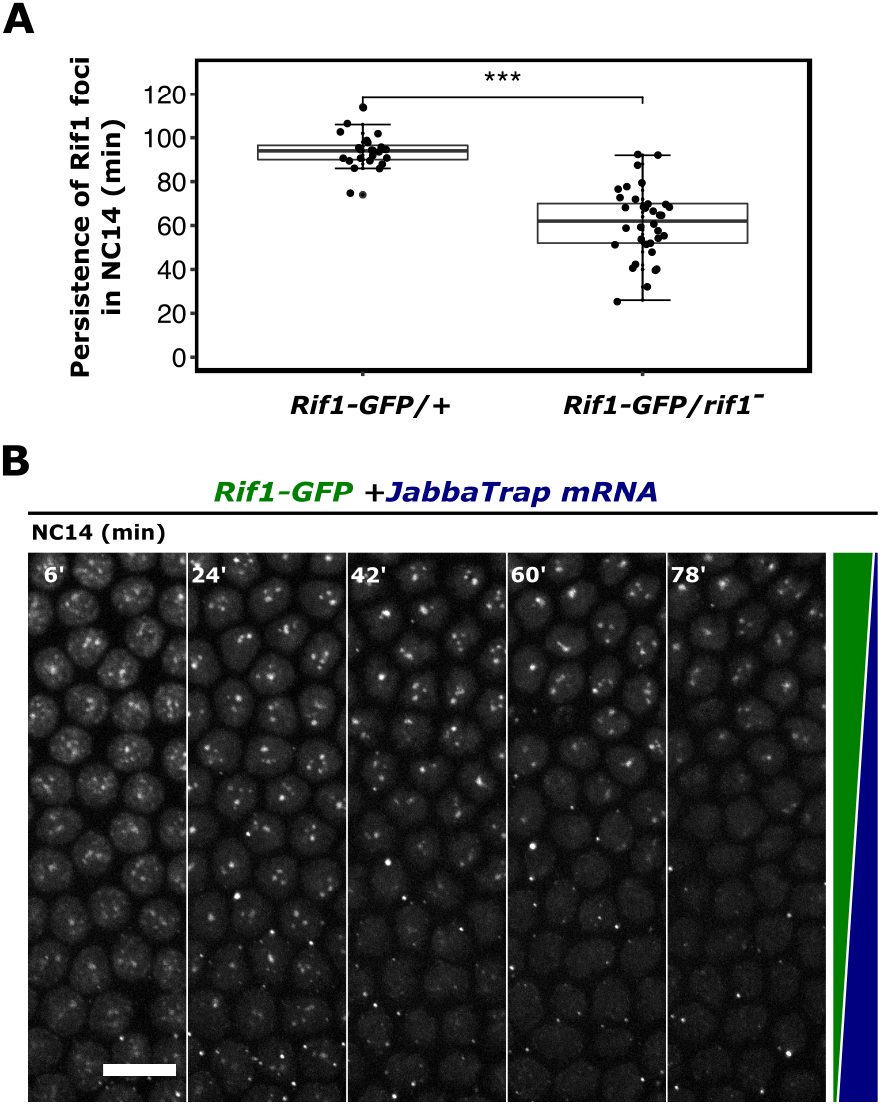
The concentration of Rif1 determines the timing of its own dissociation from satellite DNA. (A) Boxplot showing persistence of Rif1 foci during NC14 in indicated genotypes. ***p<0.001 by one-tailed t-test. Individual data points represent the results from single embryos. In each embryo, the persistence of Rif1 foci was determined as the latest time point in NC14 when Rif1 foci were visible in any nucleus in the field. (B) Timelapse imaging of Rif1-GFP during NC14 in an embryo injected with mRNA encoding JabbaTrap toward anterior pole (bottom) during the previous cycle. The JabbaTrapping depletes nuclear foci (green schematic) and creates cytoplasmic foci (blue). The start of interphase is set as time point 0’.

As a further test of the influence of Rif1 levels on the timing of Rif1 foci dispersal, we used the JabbaTrap (a fusion of the Jabba protein with an anti-GFP nanobody) to inhibit the GFP-tagged Rif1 by sequestering it at cytoplasmic lipid droplets (11). We injected mRNAs for the JabbaTrap towards one pole of Rif1-GFP embryos in NC13 and followed the persistence of nuclear Rif1 foci at the MBT by live imaging. The injection of mRNAs resulted in a gradient of JabbaTrap expression, leading to a gradient in cytoplasmic trapping of Rif1-GFP that was visualized as a gradient of cytoplasmic foci along the embryonic anterior-posterior axis (Figure 3B). An inverse gradient was seen in nuclear Rif1-GFP foci (Figure 3B). In the area close to the injection site, Rif1-GFP was largely localized to cytoplasmic foci, and fewer nuclear Rif1 foci were observed, suggesting that the initial formation of nuclear Rif1 foci is also influenced by Rif1 nuclear concentration. In these nuclei, the foci also disappeared earlier compared to those further away from the injection site with higher levels of nuclear Rif1 (Figure 3B). We conclude that the abundance of Rif1 controls the persistence of its own foci and thus the timing of late replication at satellite DNA loci.

### The balanced levels of DDK and Rif1 governs the timing of Rif1 foci dissociation

Finally, we wanted to determine whether the dosage-dependent stability of Rif1 foci is counteracted by S-phase kinases. Both Cdk1 and DDK have been shown to inhibit Rif1 before the MBT (10). However, at the MBT during NC14, Cdk1 activity is abruptly down-regulated and remains inactive until transcription of Cdc25/String as each mitotic domain triggers progression into cycle 15 (37–39). We thus hypothesized that DDK predominantly triggers the dissociation of Rif1 foci during S phase of NC14, and that inhibiting DDK activity during NC14 should delay the disappearance of Rif1 foci.

A small-molecule inhibitor of Cdc7 kinase, XL413 (referred to as Cdc7i herein), has been shown to inhibit *Drosophila* DDK activity *in vitro* (40). Previous data showed that maternal depletion of Cdc7 by RNAi severely disrupted the nuclear division cycles after fertilization, and that this disruption can be substantially rescued by the *rif1* deletion (10). If Cdc7i effectively inhibits DDK activity *in vivo*, we expect its action to parallel these Cdc7-depletion findings. We thus injected Cdc7i into wild-type embryos during mitosis 11 and followed the subsequent cell-cycle progression by live imaging of mCherry-PCNA. We observed retention of mCherry-PCNA on chromosomes well into mitosis 12 in the drug-treated wild-type embryos (Figure 4A, upper 3 and 6-min panels). Subsequent anaphase entry produced severe chromosomal bridges (Figure 4A, upper 9-min panel), suggesting a failure to complete replication. This is expected for substantial but incomplete inhibition of origin firing, leaving few forks to replicate long tracks of DNA. Importantly, while wild-type embryos injected with Cdc7i exhibited anaphase bridges immediately during the next mitosis, similarly injected *rif1* mutant embryos did not exhibit cellcycle defects and went through multiple nuclear division cycles normally (Figure 4A, lower panels; n=10 embryos). This suppression of the drug-induced phenotype is consistent with the previously documented suppression of Cdc7-RNAi-induced embryonic phenotypes by genetic loss of Rif1 function, a finding that led us to the conclusion that the level of Cdc7 required to fire origins is minor compared to the level required to inhibit Rif1 and de-repress origins (10). These results support the efficiency and specificity of the Cdc7i XL413 in reducing DDK activity in *Drosophila* embryos.

**Figure 4.**
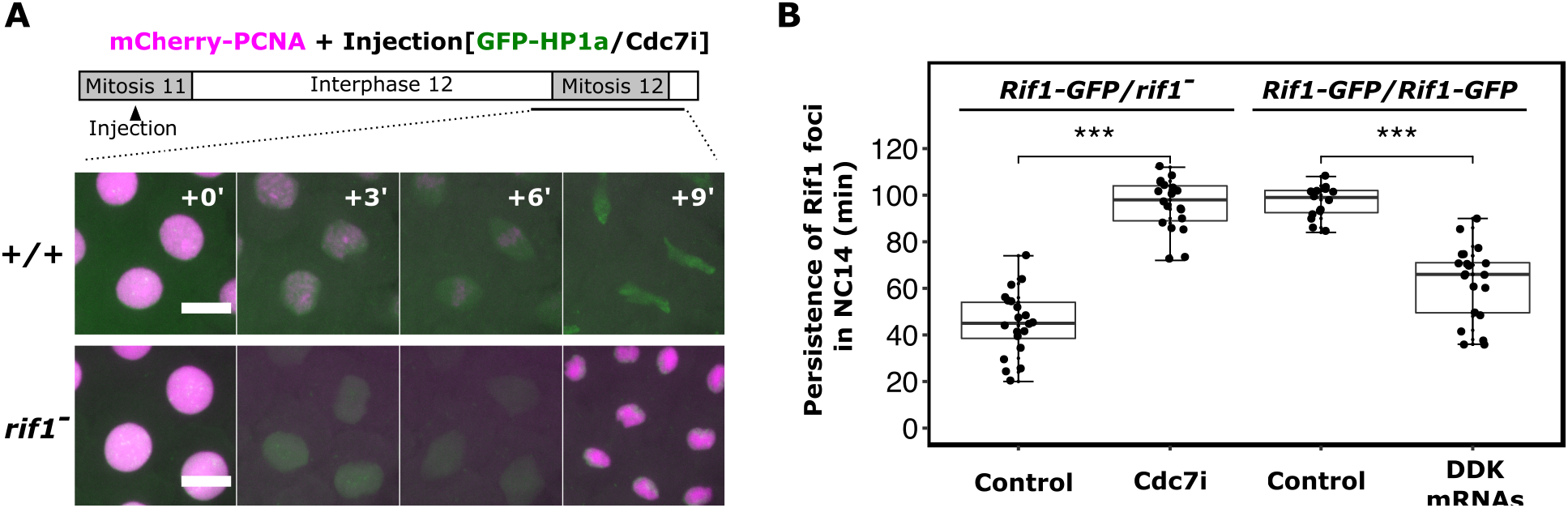
The balanced levels of Cdc7 kinase and Rif1 time the dissociation of Rif1 foci. (A) Time-lapse imaging of wild-type or *rif1^-^* embryos expressing mCherry-PCNA from a transgene and injected with XL413 (Cdc7i) along with GFP-HP1a as an injection marker. The injection was completed before interphase 12 in both experiments. (B) Boxplot showing persistence of Rif1 foci during NC14 in indicated experiments. ***p<0.001 by one-tailed t-test. Each data point represents individually scored embryo. The persistence of Rif1 foci is determined manually as the last time point in NC14 when any nucleus in the field still has visible Rif1-GFP foci.

We then assayed if abruptly reducing DDK activity during NC14 by Cdc7i injection could stabilize Rif1 foci. We used embryos with one dose of functional Rif1 (from *Rif1-GFP/rif1* mothers), because their shortened program (Figure 3A) provided a sensitized background for scoring an extension of the program. Indeed, the injection of Cdc7i greatly prolonged the persistence of Rif1 foci as compared to the control injection (Figures 4B and S4A). Notably, active replication foci marked by mCherry-PCNA exhibited two distinct waves in the Cdc7i-injected embryos. The initial foci of PCNA dispersed well before the loss of many Rif1 foci (around 40-min in Movie S1), as if replication had finished (a pseudo G2). After a long period of low PCNA nuclear intensity, PCNA again formed foci, this time at locations marked by persistent Rif1 shortly after the dispersal of Rif1 (Figure S5 and Movie S1). This late replication occurs at different times in different cells in a well-documented spatiotemporal pattern anticipating entry into mitosis 14 governed by the onset of zygotic expression of the Cdk1 activator, Cdc25/string (37, 39). Taken together with the findings that ectopic Cdc25 in early NC14 dissociates Rif1 and shortens S phase (10, 41), we suggest that endogenous Cdc25 expression and the associated rise in Cdk1 activity toward the end of interphase trigger Rif1 release and allow the completion of S phase. Thus, this Cdk1-mediated dissociation of Rif1 serves as a fail-safe program to prevent mitotic catastrophe.

In a final test of the contribution of DDK to the timing of Rif1 dissociation, we overexpressed DDK in the homozygous Rif1-GFP embryos by injecting mRNAs before NC13, allowing DDK accumulation by NC14 when we examined the effects on Rif1 foci. The disappearance of Rif1 foci in the Rif-GFP homozygote was indeed accelerated (Figures 4B and S4B). This gain-of-function experiment shows that DDK can antagonize the dose-dependent stability of Rif1 foci to control the timing of late replication, while the inhibitor experiment shows that its activity normally contributes to the timing of Rif1 foci dissociation and the timing of replication of satellite sequences.

## DISCUSSION

In this study, we have investigated the mechanisms that control the timing of Rif1 foci dispersal from satellite sequences, which dictates the onset of late replication in the MBT embryo. We demonstrate that Rif1-recruited PP1 mediates self-stabilization of Rif1 hubs, while the S-phase kinase DDK opposes PP1 action and triggers the dispersal of Rif1 hubs. We propose a model in which the firing of late origins is primarily controlled by a de-repression step upstream of the activation of the pre-RC. In this model, hubs of Rif1 create domains of locally high PP1 that prevent kinase activation of underlying pre-RCs. However, a changing balance of local phosphatase and kinase levels leads to the abrupt destabilization of different Rif1 hubs at distinct times (Figure 5). This alleviates PP1 inhibition of hub-associated origins at specific times to trigger replication of the different satellites at different times. While this simple model appears sufficient to explain the late replication at its initial onset in the early *Drosophila* embryo, numerous other factors impact the replication program at later stages when chromatin acquires more complex features (42). Nonetheless, as we discuss below, the simplicity of the process in this biological context offers some insights into the more enigmatic aspects of late replication, and perhaps suggests a flexible regulatory paradigm that might be used in diverse contexts.

**Figure 5.**
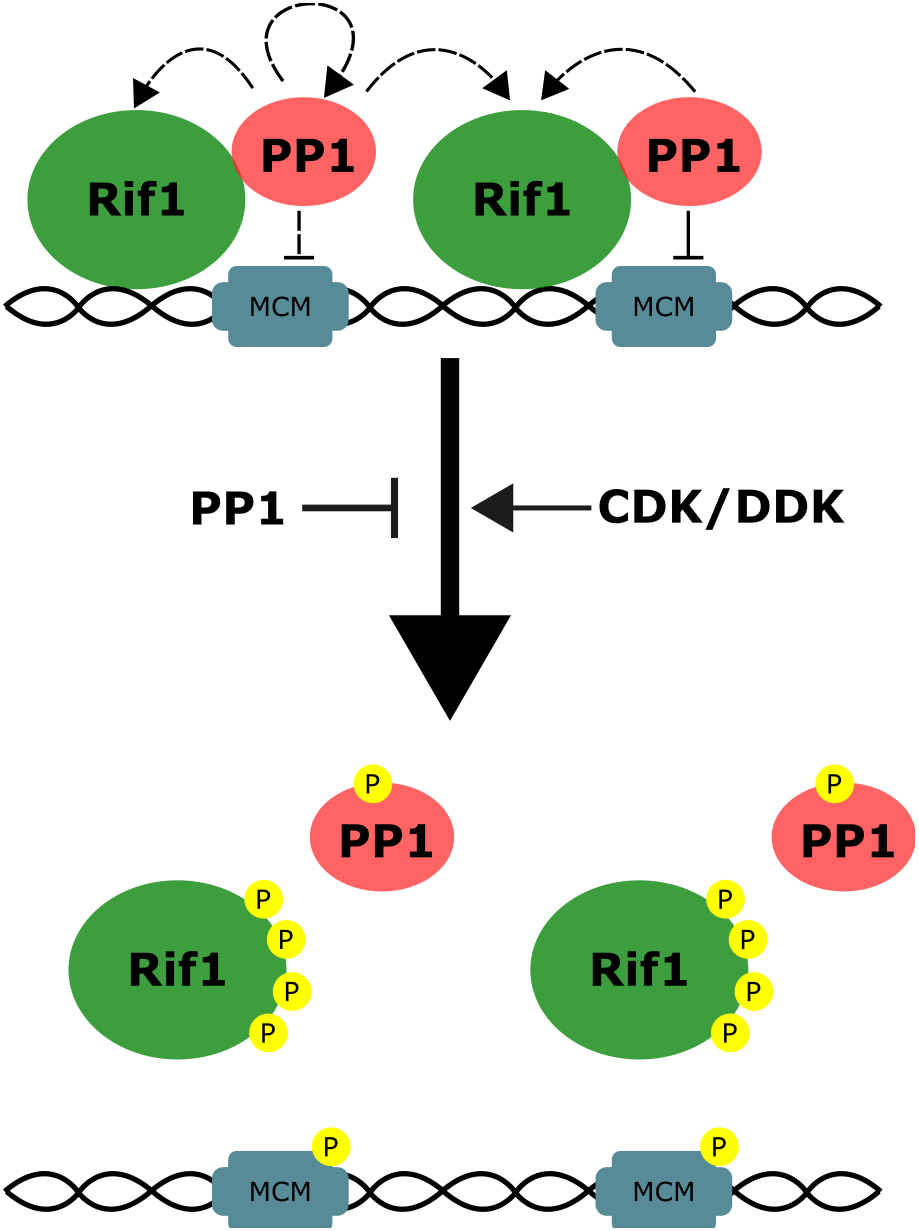
A model for the multiple actions of PP1 in stabilizing Rif1 hubs.

While the mechanism is unknown, it has long been clear that large domains of the genome behave as timing units, and that the numerous origins within such domains fire coordinately if not synchronously (1). The “hub” model of late replication control in the early embryo can explain how the firing of numerous origins within megabase pairs of satellite sequences can be coordinated in late S phase. Each Rif1 hub is associated with a locus of repetitive satellite sequence (10). Coordinated dispersal of a Rif1 hub will convert the subnuclear compartment from one restricting kinase actions to a permissive one, allowing the activation of pre-RCs throughout the associated chromatin domain. It was previously unclear what leads to the coordinated dispersal of these large hubs. Here we show that a mutant Rif1 that is deficient in binding PP1 cannot form stable hubs on its own, but it joins wild-type Rif1 in semi-stable hubs. Importantly, the mutant and wild-type Rif1 disperse together, showing that they respond equally to the property of the domain. We suggest that Rif1-bound PP1 can act in *trans* to stabilize nearby Rif1-PP1 and that the propagation of this action coordinates the behavior of Rif1 across the entire hub.

The contribution of PP1 to the self-stabilization of Rif1 hubs might be mediated by feedback at multiple levels (Figure 5): 1. PP1 might activate itself by removing inhibitory phosphorylation catalyzed by Cdk1 (43); 2. It could reverse Cdk1/DDK-mediated phosphorylations of Rif1 that disrupt PP1-recruitment (24, 25, 44); 3. It could reverse phosphorylations of Rif1 that disrupt Rif1 chromatin association (10); 4. Or in a circuitous pathway, if the firing of origins were to promote Rif1 dissociation, PP1-dependent suppression of origin firing would stabilize the hubs. Any or all the above actions could reinforce the stability of Rif1-PP1 hubs, perhaps making different contributions in different situations and different organisms. However, regardless of the feedback route, a local dominance of PP1 will stabilize the Rif1 hubs, and rising kinase activity could erode this dominance of PP1. Upon reaching a tipping point, the local PP1 would no longer successfully stabilize the Rif1 hub, and S-phase kinases would then trigger complete dispersion and allow replication of the underlying chromatin.

A potential ability of origin firing to feedback and destabilize Rif1 hubs might explain observations in other organisms suggesting that the level of a variety of replication initiation factors can influence replication timing. For example, overexpression of four replication factors including a DDK subunit in the *Xenopus* embryo shortens the S phase at the MBT (45). While this has been interpreted as evidence for governance of replication timing by limitation for these factors, over-production of all of these factors might override Rif1 suppression of pre-RC activation, and the resulting activation could destabilize Rif1 hubs.

Importantly, the replication defects resulting from Cdc7 knockdown (10) or inhibition of Cdc7 (Figure 4), are suppressed in a *Rif1* null mutant background. This shows that the level of DDK activity required to reverse or override Rif1 suppression of pre-RC activation is greater than the level needed for direct pre-RC activation. Thus, in a scenario in which rising levels of DDK during S-phase 14 act as a timer, genomic domains associated with Rif1 hubs would fail to replicate until DDK reached the high level required to destabilize the hub. This argues that replication timing depends on the threshold for derepression of the domain rather than on distinct thresholds for firing individual pre-RCs. We therefore suggest that the timing of late replication is governed at the level of the upstream derepression step in *Drosophila* embryos, in contrast to the model proposed for other organisms according to which activation of pre-RCs are directly limited by availability of DDK and other replication factors (46, 47). To produce the distinct temporal program of replication of different satellites, our model requires domain-specific distinctions in the threshold for hub dispersal. Different satellite loci that are composed of a common repeat sequence replicate at the same time, while satellites composed of different sequences replicate at distinct times. This leads us to propose that the sequence of repeats influences, likely indirectly, the threshold for Rif1 hub dispersal and the timing of replication.

The possible generality of the circuitry we have defined in the cycle 14 *Drosophila* embryo can be considered in various ways. Focusing directly on Rif1 involvement in late replication, it is clear that Rif1 does not bare full responsibility for late replication at other stages. Nonetheless, a dosage-dependent function of Rif1 in controlling replication timing is also observed in *Drosophila* follicle cells during their mitotic cycles (16). Furthermore, in mammalian cells, ChIP-seq and microscopy showed that Rif1 interacts with large late-replicating domains but, as we see in cycle 14 embryos, is absent once onset of replication of the underlying chromatin is detected (12, 21). We suggest that the mechanism we have described will be one of multiple contributors to replication timing control in other biological contexts, and it is likely to be the major mode of replication timing in the rapid cycles of externally developing animal embryos.

Rif1 has other regulatory roles beyond timing control of pre-RC activation. In the follicle cells of *Drosophila* egg chambers, Rif1 is recruited to specialized replication forks during chorion gene amplification where it suppresses fork progression (34). While this action of Rif1 is dependent on its ability to associate with PP1, other possible parallels to the mechanism we describe here are not evident. Rif1 also regulates biological processes beyond replication. It is recruited to regions of DNA damage in mammals as well as to the telomeres in yeast where it has regulatory roles involving distinct interactions (48, 49). Thus, Rif1 recruitment appears to trigger alternative regulatory pathways in different circumstances.

Despite the evident diversity of biological regulation, the capacity of Rif1 to form local membraneless compartments dominated by phosphatase and to abruptly dissolve in response to kinase levels might be an example of a group of flexible regulatory strategies. Many important regulatory events, such as phosphorylation, acetylation, and ubiquitination, are countered by reverse reactions. Various processes, notably the formation of liquid-like condensates, promote local accumulation of proteins. Accumulations of proteins that promote or oppose regulatory modifications could control major regulatory pathways. Furthermore, since protein accumulations could be stabilized or destabilized by the modifications they regulate, a feedback mechanism could control the formation and destabilization of a compartment to give precise spatiotemporal control, as exemplified by the behavior of the Rif1 hubs in the cycle 14 *Drosophila* embryo.

## MATERIALS AND METHODS

### Fly stocks

All *Drosophila melanogaster* stocks were grown on standard cornmeal-yeast medium. Strains used in this study are as follows: the *w^1118^* Canton-S as wild type, *Rif1-EGFP, rif1^-^*, *mCherry-PCNA^attP-9A^* (10), and *Rif1^pp1^-EGFP* (this study).

### CRISPR-Cas9 mutagenesis

To generate *Rif1^pp1^-EGFP*, we first introduced *vas-Cas9* (Bloomington #51324) into the *Rif1-EGFP* line. The sgRNA targeting 3’ end of PP1-docking motif was cloned into pU6-BbsI-chiRNA and co-injected with single-stranded oligo DNA donor carrying Rif1^pp1^ mutation into the above line. Surviving adults were crossed to CyO balancer strain and screened by PCR and Sanger sequencing for successful mutagenesis. *vas-Cas9* was removed by further outcross with wild type. The injection was performed by Rainbow Transgenic Flies, Camarillo, CA.

### In vitro transcription of mRNA

All DNA templates were cloned into a vector backbone downstream of T7 promoter sequence. The Rif1^15A^ and Rif1^15A.pp1^ cDNAs were modified from previously made Rif1^15A^-EGFP plasmid using Gibson assembly for mutagenesis. Full-length *Cdc7* and N-terminal *Chiffon* cDNAs encoding Dbf4 homolog (40) were amplified from a general cDNA library prepared from embryos as described (31). Linear DNA templates were obtained by either restriction enzyme digestion or PCR amplification, and mRNA was produced by CellScript T7 mScript Standard mRNA Production System and resuspended in ddH2O (31).

### Embryo collection and microinjection

All heterozygous females were generated by crossing *Rif1-EGFP* or *Rif1^pp1^-EGFP* with either wild type or *rif1^-^*. F1 females were crossed with siblings, and their embryos were collected for experiments. Embryos were collected on grape agar plates, aged at 25°C, washed, and dechorionated in 50% bleach for 2 minutes. Embryos were then aligned, glued to glass coverslips, and covered in halocarbon oil for live imaging.

Microinjection was performed as described (31). Embryos were handled as above but were desiccated for 8-12 minutes before covering with halocarbon oil. Recombinant mCherry-PCNA and GFP-HP1 proteins were purified and injected as described previously (4). All mRNAs except for JabbaTrap were injected at ~750 ng/μl. To avoid sequestering all Rif1-GFP proteins, JabbaTrap mRNA was injected at 75 ng/μl. The Cdc7 inhibitor XL413 (Sigma, #SML1401) was injected at 0.5 mg/ml in 40 mM HEPES, pH 7.4, and 150 mM KCl with or without purified GFP-HP1 proteins.

### Microscopy

Live imaging was performed on an Olympus IX70 microscope equipped with PerkinElmer Ultraview Vox confocal system or a Leica DM 1RB inverted microscope equipped with a Yokogawa CSU10 spinning disk confocal unit. All experiments were performed at room temperature with 63x or 100x oil objective. 11 Z-stacks each 1 μm apart spanning the apical part of the nuclei were recorded every 30-120 seconds, depending on the duration of time-lapse. GFP and RFP proteins were excited using a 488 nm and a 561 nm laser respectively. Data were acquired using Volocity 6 software (Quorum Technologies). Because nuclei move slightly inward into the interior of the embryos during cellularization in NC14, focal planes were manually adjusted every 15-30 minutes between time points when imaging embryos in NC14. Images being compared were acquired and processed using identical settings unless otherwise noted. Maximal projection of Z-stacks was used for analysis in FIJI and presentation.

## ACKNOWLEDGEMENTS

We thank members of the O’Farrell lab, especially Pei-I Tsai and Ekaterina Korotkevich, and our colleague Mustafa Aydogan for critical reading and helpful discussion. This work is supported by National Institutes of Health grant GM037193 (to P.H.O).

## AUTHOR CONTRIBUTIONS

C.-Y.C. and P.O’F conceptualized the study. C.-Y.C. and C.S. performed the experiments. C.-Y.C. analyzed the data. C.-Y.C. and P.O’F wrote the manuscript. C.S. edited the manuscript.

---

**Figure S1.**
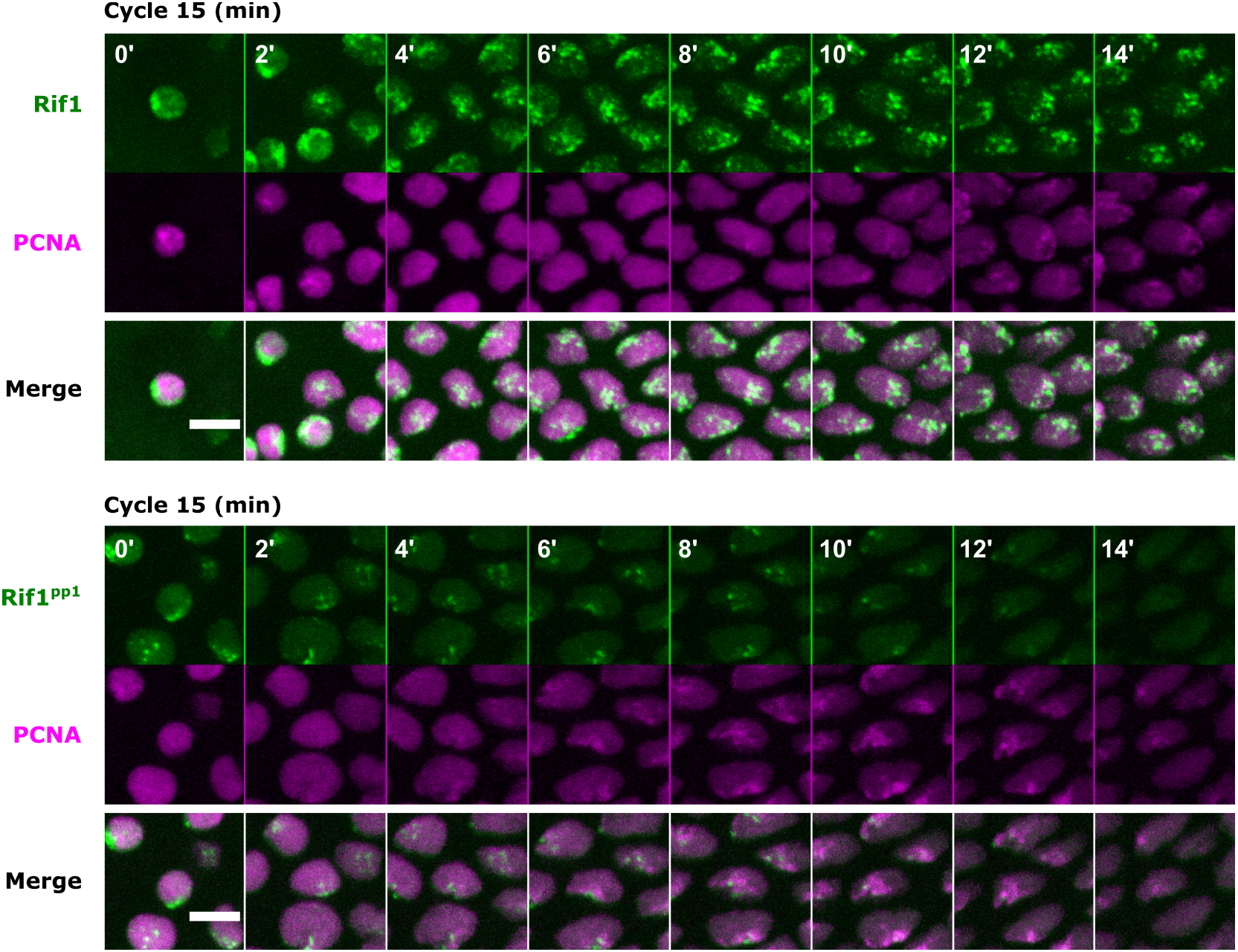
Additional analyses related to Figure 1. Time-lapse imaging of GFP-tagged Rif1 or Rif1^pp1^ along with mCherry-PCNA during cycle 15. The start of interphase is referred to as time point 0’. Scale bar, 6 μm.

**Figure S2.**
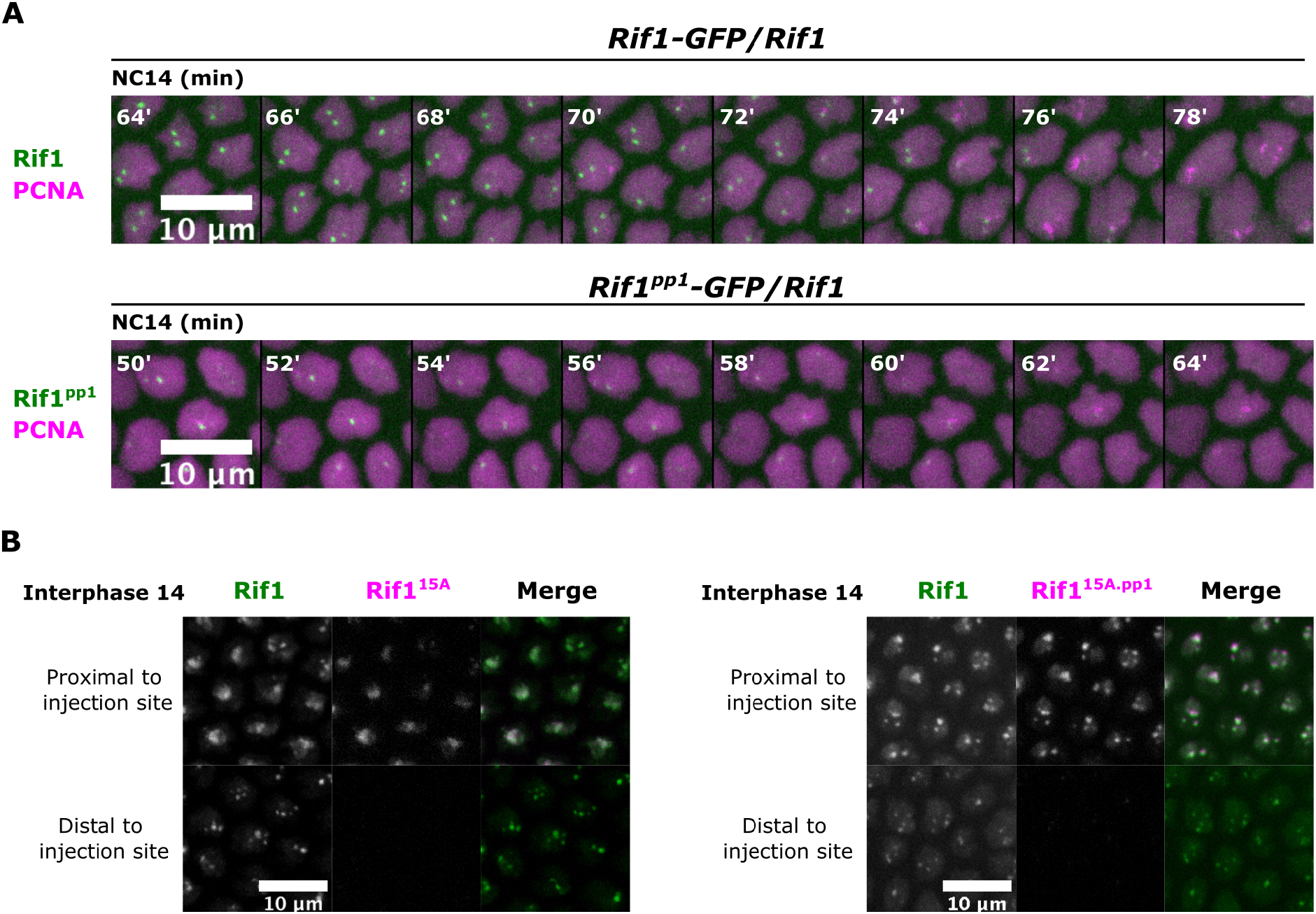
Additional analyses related to Figure 2. (A) Time-lapse imaging of embryos from females of indicated genotypes during late NC14 when Rif1-GFP foci were dissociating. The start of interphase 14 is set as time point 0’. (B) Snapshots from time-lapse movies following Rif1-GFP embryos injected with mRNAs encoding Rif1^15A^ mutants tagged with mScarlet-I. Nuclei proximal or distal to injection sites about 1 hour in S phase 14 are shown.

**Figure S3.**
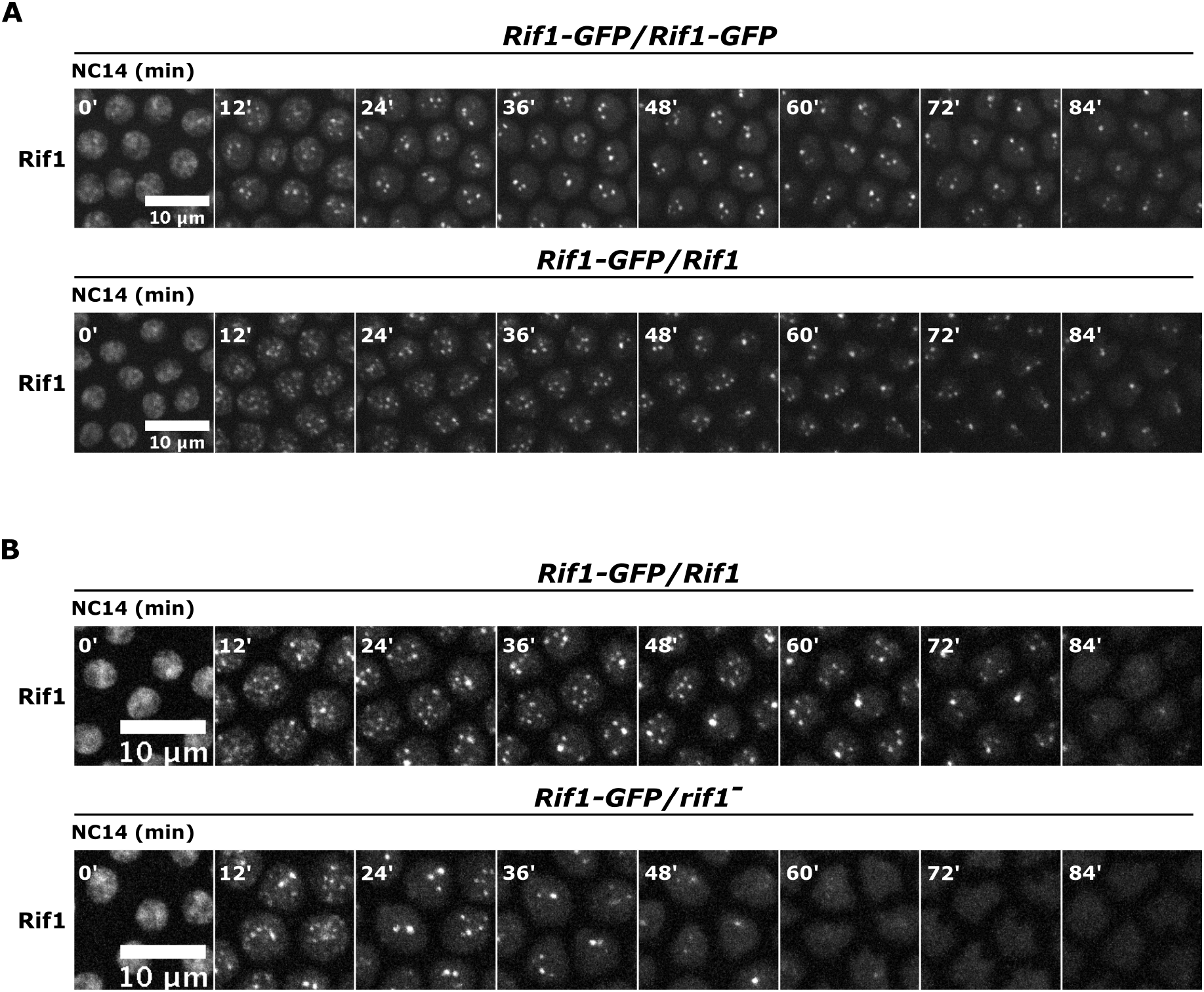
Additional analyses related to Figure 3. (A) Time-lapse imaging of Rif1-GFP in embryos from females carrying one or two copies of GFP-tagged Rif1 during cycle 14. (B) Time-lapse imaging of Rif1-GFP in embryos from females trans-heterozygous for Rif1-GFP with either wild-type *Rif1* or a *rif1* deletion.

**Figure S4.**
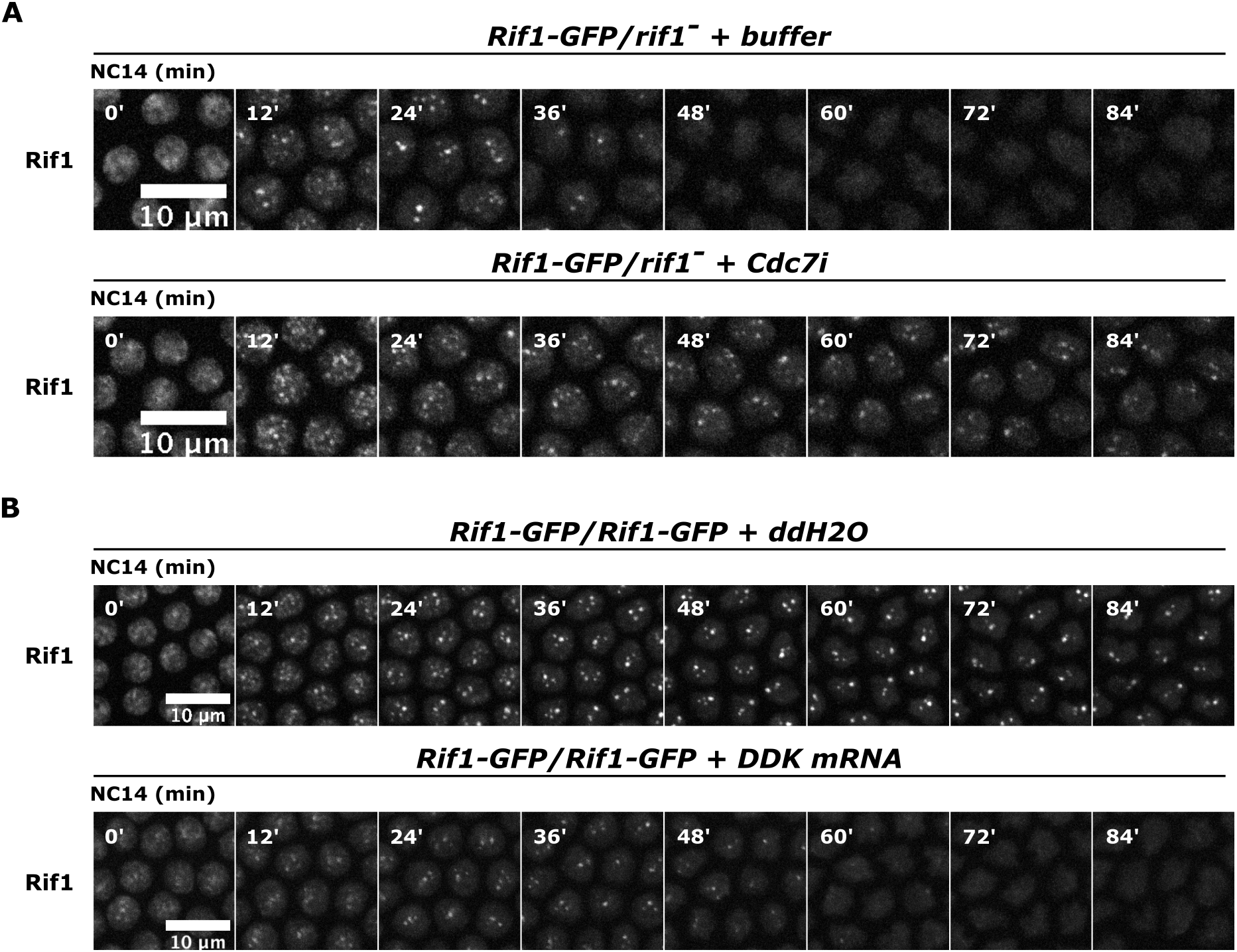
Additional analyses related to Figure 4. (A) Time-lapse imaging of Rif1-GFP in cycle 14 embryos from *Rif1-GFP/rif1* heterozygous females and injected with buffer or Cdc7i. The injection of Cdc7i was performed during mitosis 13. (B) Time-lapse imaging of Rif1-GFP in cycle 14 embryos from *Rif1-GFP* homozygous females and injected with control water or mRNAs encoding DDK subunits. The injections were performed before cycle 13.

**Figure S5.**
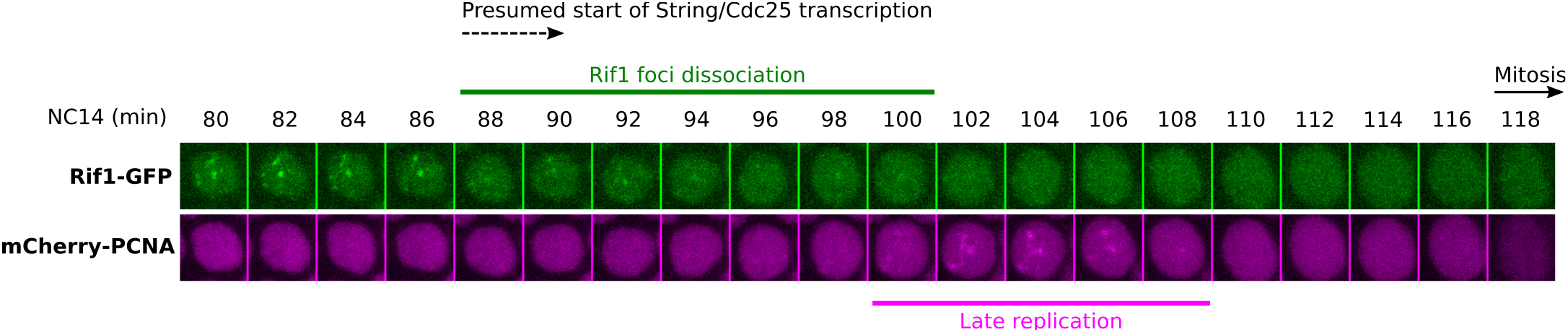
Partial inhibition of Cdc7 results in a second wave of replication controlled by a “fail-safe” program associated with the mitotic entry program. Snapshots captured from Movie S1 focusing on a nucleus in the embryo injected with Cdc7i during late NC14. The time of String/Cdc25 expression is taken from measurements of the lag between the first detectable zygotic transcription and onset of mitosis (37).

**Movie S1. Inhibition of Cdc7 kinase activity stabilizes Rif1 foci during cycle 14.** Time-lapse imaging of Rif1-GFP and mCherry-PCNA in a cycle 14 embryo from heterozygous *Rif1-GFP/rif1* females and injected with either buffer or Cdc7i. Areas near the cephalic furrow are shown with anterior pole toward the left.

